# Cell-Type Specific Molecular and Functional Consequences of TDP-43 Loss-of-Function in Human Induced Neurons

**DOI:** 10.64898/2026.01.26.700683

**Authors:** Vera Gysin Filippa, Karsten Bach, Vitaliy Kolodyazhniy, Lars Joenson, Marcos Romualdo Costa

## Abstract

Amyotrophic Lateral Sclerosis (ALS) is a devastating neurodegenerative disorder characterized by the cytoplasmic aggregation and nuclear depletion of the TDP-43 protein. The latter impairs TDP-43 function as an RNA-binding protein and compromises the repression of cryptic splicing events, affecting both glutamatergic upper motor neurons and cholinergic lower motor neurons. This study systematically investigated the molecular and functional consequences of TDP-43 knockdown in human induced pluripotent stem cell (hiPSC)-derived glutamatergic neurons (iGNs) and cholinergic motor neurons (iMNs) using antisense oligonucleotides. The results demonstrated that TDP-43 loss elicits widespread, cell-type-specific changes in gene expression and alternative splicing. Notably, a shared subset of ALS-associated targets, including STMN2 and UNC13A, were consistently downregulated and mis-spliced across both neuronal subtypes. Functionally, Microelectrode Array (MEA) electrophysiology recordings revealed that TDP-43 knockdown induces a hyperexcitable phenotype in both neuronal populations, though they exhibited distinct network patterns: iGNs displayed synchronized bursting and significant shifts in overall electrophysiological profiles, while iMNs showed asynchronous firing. Furthermore, the inclusion of astrocytes in co-culture models expanded the repertoire of detectable cryptic splicing, including an event in HDGFL2 previously identified in patient cerebrospinal fluid. Despite these profound molecular and functional deficits, TDP-43 depletion did not impact neuronal viability or increase susceptibility to glutamate-induced excitotoxicity. These findings validate hiPSC-derived iGNs and iMNs as relevant models for ALS and highlight the critical necessity of considering cell-type specificity when elucidating pathogenesis and developing targeted therapies.

## Introduction

Amyotrophic Lateral Sclerosis (ALS) is a progressive and fatal neurodegenerative disorder defined by the selective loss of motor neurons in the brain and spinal cord, affecting both upper (glutamatergic) and lower (cholinergic) motor neurons. While the etiology is complex and multifactorial, a key pathological hallmark present in up to 97% of ALS cases is the cytoplasmic mislocalization and aggregation of the TAR DNA-binding protein 43 (TDP-43), leading to its functional depletion in the nucleus ^1^. This loss of nuclear TDP-43 function is hypothesized to be a significant driver of disease pathogenesis.

TDP-43 is an essential RNA-binding protein, primarily localized to the nucleus, where it performs crucial roles in various aspects of RNA processing, including transcription, splicing, mRNA transport, and translational regulation ^2,3^. A key function is its role in the repression of cryptic splicing variants through binding to specific intronic regions of pre-mRNA ^4,5^. Consequently, the loss-of-function associated with nuclear TDP-43 depletion results in the aberrant inclusion of normally suppressed cryptic exons (CEs) in the transcripts of numerous target genes. These splicing alterations can lead to the production of non-functional or truncated proteins, or altered protein expression levels, ultimately contributing to the progressive neuronal dysfunction and degeneration observed in ALS ^6^.

Understanding the precise splicing defects caused by TDP-43 loss-of-function and their downstream consequences is critical for elucidating ALS pathogenesis and identifying potential therapeutic targets. Reduced expression of TDP-43, for instance, has been shown to decrease microtubule outgrowth in motor neurons through the premature polyadenylation and subsequent loss of Stathmin-2 (STMN2) transcripts ^7,8^. Similarly, the TDP-43 depletion-associated loss of Unc-13 homolog A (UNC13A) has been linked to alterations in electrophysiological properties, particularly synaptic function ^9^.

Cryptic splicing events are not unique to TDP-43 proteinopathy but have been consistently shown in a wide range of ALS and frontotemporal dementia (FTD) patient data, as well as in rodent and in vitro models ^6,10^. Recently, studies have demonstrated that these TDP-43-related splicing changes may exhibit tissue or cell-type-specific patterns. For example, a study by Newton and Friedman (2024) identified distinct spinal and cortical splicing signatures across ALS/FTD tissue datasets, further describing a specific in vitro signature unique to TDP-43 knockout cell models ^10^. Similarly, Jeong and colleagues identified specific subsets of splicing changes by cell-type-specific TDP-43 knockout in mouse muscle cells or neurons ^11^. Yet, a gap remains in understanding whether similar genes are regulated by TDP-43 in different neuronal subtypes, such as the glutamatergic and cholinergic neurons that are critically vulnerable in human ALS. Addressing these cell-type-specific molecular consequences is vital for validating in vitro models relevant to both upper and lower motor neuron degeneration.

Furthermore, neuronal hyperexcitability—characterized by increased neuronal firing and heightened susceptibility to synaptic input—has been repeatedly described in both ALS patients and mouse disease models ^12–15^. This hyperexcitable phenotype has also been observed in ALS in vitro models, including patient-derived iPSC models ^16^. The clinical relevance of this phenomenon is highlighted by Riluzole, the first FDA-approved drug for ALS, which modulates glutamatergic neurotransmission to reduce excitotoxicity and neuronal hyperexcitability, offering a modest extension of survival ^17^. Understanding how TDP-43 loss-of-function affects neuronal electrical properties and the specific molecular alterations that underpin this functional change may, therefore, pinpoint new potential therapeutic targets.

Here, we systematically investigated the molecular and functional consequences of TDP-43 knockdown in two different subtypes of human induced neurons (hiNs): hiPSC-derived glutamatergic neurons (iGNs) and hiPSC-derived cholinergic motor neurons (iMNs). Our study combined transcriptomic analysis of gene expression and alternative splicing with microelectrode array (MEA) electrophysiology and neurodegeneration assays to reveal cell-type-specific molecular profiles and functional deficits resulting from TDP-43 loss-of-function.

## Results

### TDP-43 knockdown elicits cell-type specific changes in gene expression

To investigate the effects of TDP-43 loss-of-function in neuronal subtypes selectively vulnerable in ALS, we utilized human induced pluripotent stem cell (hiPSC)-derived glutamatergic neurons (iGNs), modeling upper motor neurons, and cholinergic motor neurons (iMNs), modeling lower motor neurons. To achieve a controlled reduction of TDP-43, we optimized an Antisense Oligonucleotide (ASO)-mediated knockdown strategy (Figure 1A). Using three doses of a TDP-43-targeting ASO in iGN and iMN cultures at 21 days in vitro (DIV), we determined that a 5 μM dose was sufficient to achieve approximately a 50% reduction in both TDP-43 protein and TARDBP mRNA levels (Figures 1C, D). This dose was selected for all subsequent experiments. After five weeks in culture, both iGNs and iMNs displayed long, complex processes, as visualized by staining for the microtubule-associated protein MAP2 and the pan-neuronal marker HuC/D (Figure 1B). Immunocytochemistry (ICC) confirmed that endogenous TDP-43 was primarily nuclear and that treatment with the TDP-43-ASO successfully reduced nuclear TDP-43 protein by approximately 50% compared to the phosphate-buffered saline (PBS) and non-targeting (NT) ASO controls (Figures 1E, F).

**Figure 1:**
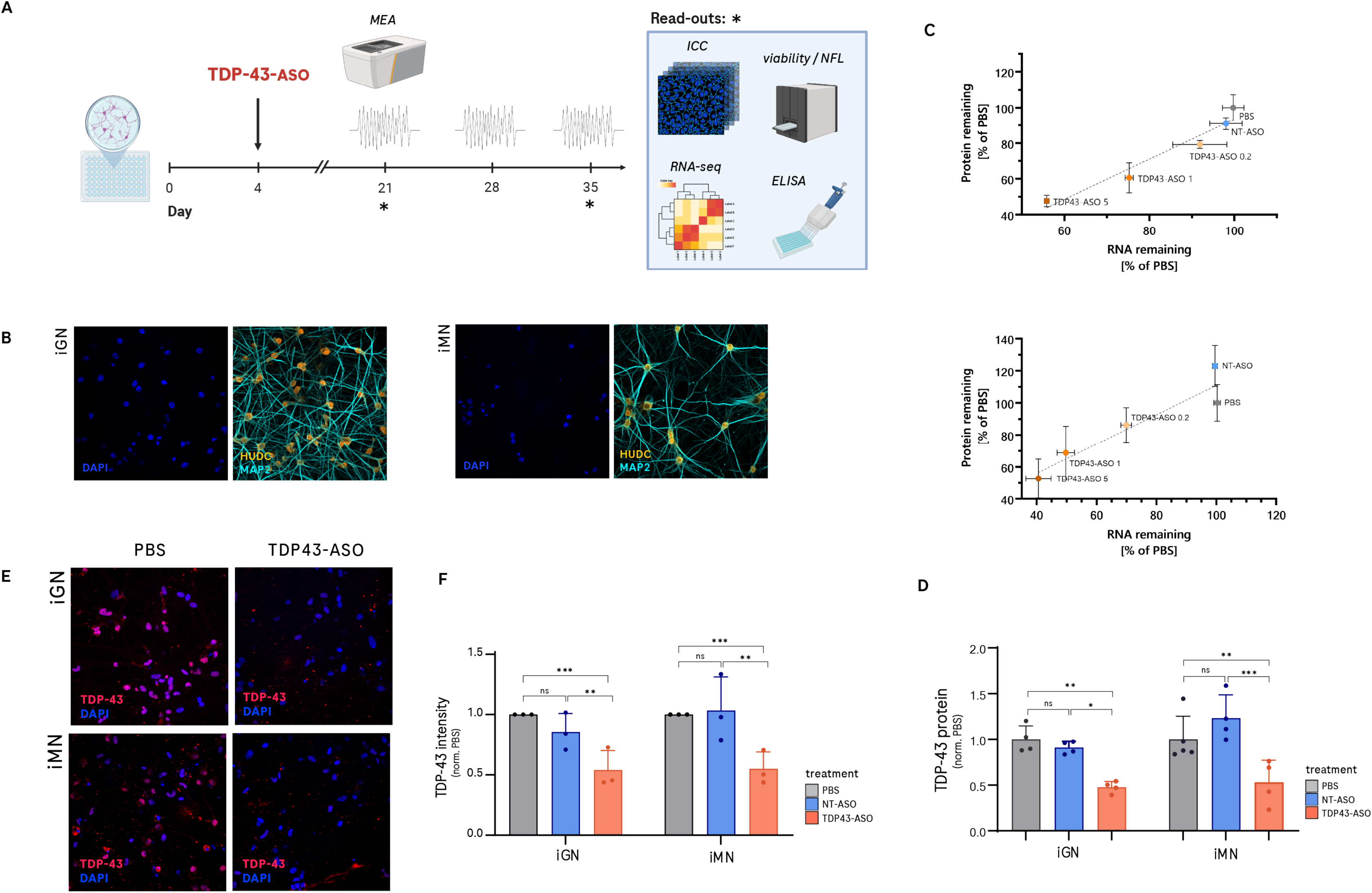
Efficient TDP-43 knockdown using an antisense oligonucleotide (ASO) in iGN and iMN cells. (A) Schematic showing the experimental timeline for cell culture, the TDP-43-ASO treatment, and assays performed, including microelectrode array (MEA), immunocytochemistry (ICC), and RNA-sequencing (RNA-seq), ELISA and viability and NfL measures. (B) Representative ICC images of iGN and iMN cells stained for HUDC, MAP2, and DAPI. (C) Correlation of TDP43 protein remaining (measured by ELISA) and TARDBP RNA at 21 div at three TDP43-targeting ASO concentrations (5uM, 1uM, 0.2uM) and two control conditions (PBS, non-targeting ASO). (D) Quantification of TDP-43 protein levels normalized to the mean of PBS control (n=4), one-way ANOVA with Dunnett’s post-hoc (p** < 0.001, p** < 0.01, p* < 0.05). (E) Representative ICC images showing the effect of the TDP-43-ASO on nuclear TDP-43 in iGN and iMN cells compared to a PBS control. Cells were stained for TDP-43 and DAPI. (F) Quantification of the nuclear TDP-43 intensity from the ICC images, normalized to the mean of PBS control (n=3, independent experiments), one-way ANOVA with Dunnett’s post-hoc (p** < 0.01, p* < 0.05).

To characterize the molecular consequences of reduced TDP-43 expression, we performed bulk RNA sequencing on iGNs and iMNs after five weeks of differentiation. We first confirmed a significant reduction of TARDBP mRNA levels upon ASO treatment compared to non-targeting (NT) ASO (Figures 2A, B, E). Next, we identified 92 differentially expressed genes (DEGs) (log_2_FC>1 and adjusted P value<0.05) in TDP-43 knockdown iMNs, and 52 DEGs in knockdown iGNs, both compared to their respective NT ASO controls (Figures 2A, B, Table S1 and S2). A direct comparison of the gene expression changes revealed a significant degree of cell-type specificity. While most genes were uniquely affected in each cell type, only a subset of 33 genes (8 upregulated, 25 downregulated) showed consistent expression changes in both iGNs and iMNs (Figures 2C, D). This overlapping set, however, included several critical, previously described TDP-43 targets ^7,18^. Notably, STMN2, UNC13A, CAMK2B, ELAVL3 and ATG4B were all significantly downregulated in both iGNs and iMNs (Figure 2E).

**Figure 2:**
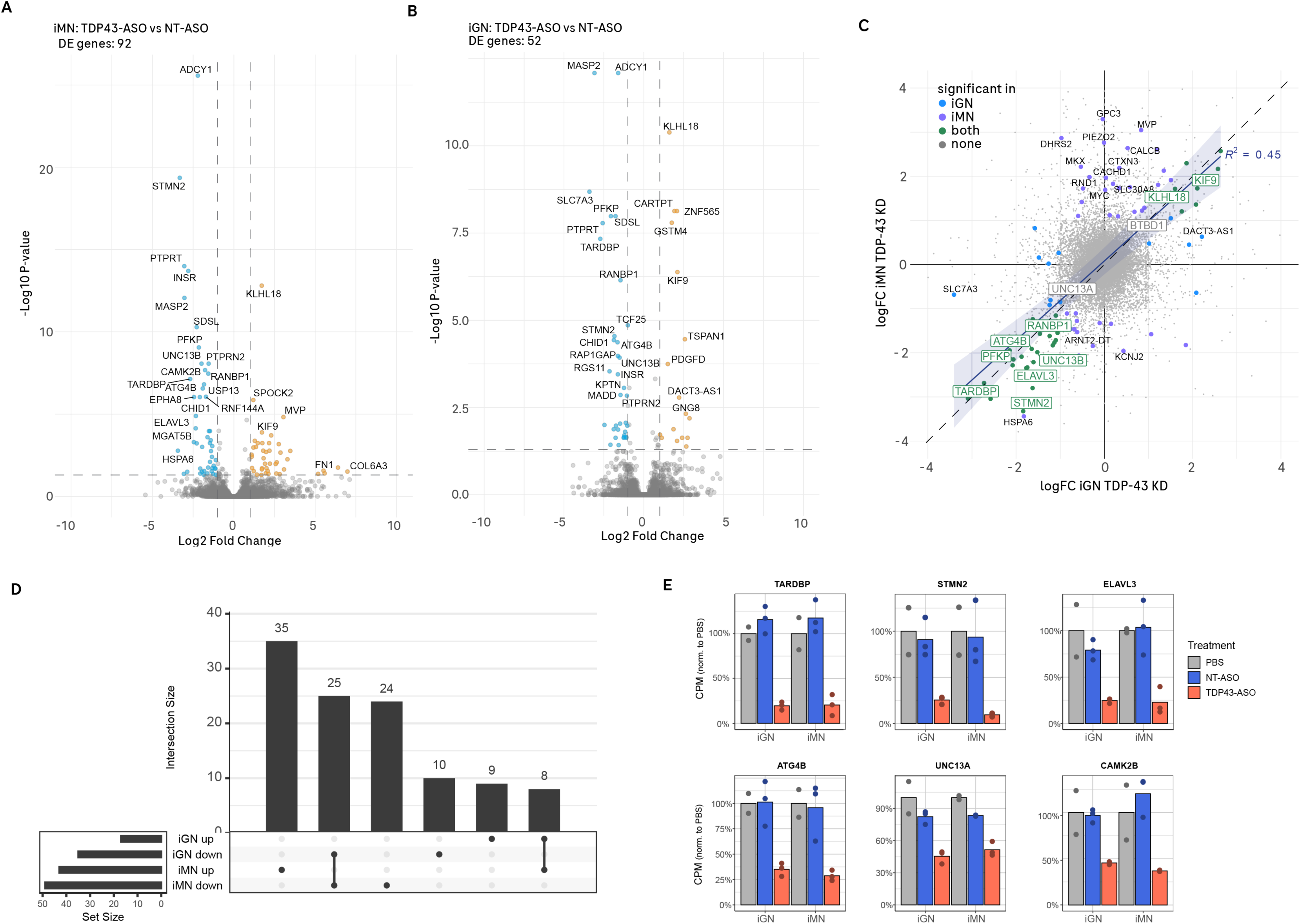
Similar and disparate gene expression changes in iGNs and iMNs following TDP-43 knockdown. (A) Volcano plot showing the differentially expressed (DE) genes in iMN cells treated with TDP-43-ASO compared to the NT-ASO control. A total of 92 DE genes were identified. (B) Volcano plot showing the DE genes in iGN cells treated with TDP-43-ASO versus the NT-ASO control. 52 genes were identified. (C) Scatter plot comparing the log2 fold change (logFC) in iGN and iMN cells after TDP-43-ASO treatment. Highlighted genes are significantly affected (FDR < 0.05) in iGN cells (blue), iMN cells (violet), or both (green). (D) UpSet plot showing the intersection of upregulated and downregulated genes in the iGN and iMN cell lines. (E) Bar graphs showing the expression levels, measured in counts per million (CPM), normalized to the PBS control for genes: TARDBP, STMN2, ELAVL3, ATG4B, UNC13A, and CAMK2B (n=2 for PBS, n=3 for NT-ASO and TDP-43 ASO).

### Splicing Alterations in Neurons and Neuron-Astrocyte Co-cultures Recapitulates Patient Data

To probe whether the gene expression changes observed in TDP-43 knockdown iMNs and iGNs were linked to altered pre-mRNA processing, we analyzed changes in alternative splicing variants using the variant frequency (VF) metric, defined as the ratio of reads for a specific variant to the total reads for that gene event (Figure 3A). We identified a substantial number of significant splicing variants down or upregulated following TDP-43 knockdown: 192 in iGNs and 93 in iMNs (Figures 3B, C, Table S3 and S4). Many of these variants represented the inclusion of normally repressed CEs. Consistent with the gene expression data, many previously identified disease-relevant splicing variants were affected in both cell lines, showing a strong correlation in their VF changes between iGN and iMN responses (Figure 3D). However, we also observed splicing variants that were not associated with a subsequent decrease in gene expression, and vice-versa (Figures 3E, F). This discrepancy suggests that other post-transcriptional mechanisms, such as nonsense-mediated decay (NMD) of the aberrant transcripts, may be at play, preventing the detection of a full-length transcript ^5^.

**Figure 3:**
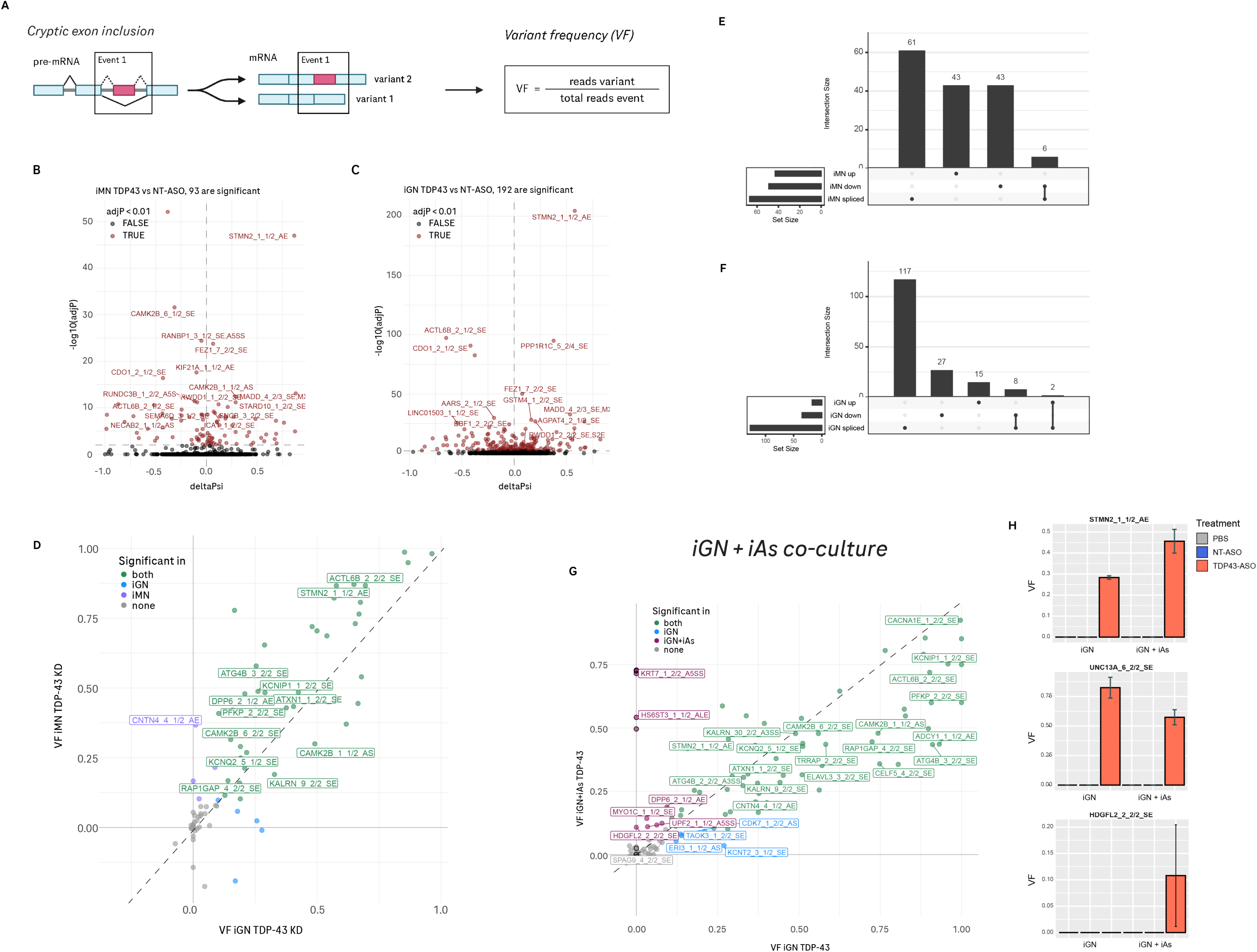
Identification of TDP-43 dependent splicing changes following TDP-43 knockdown in iNs and iAs. (A) Schematic diagram illustrating the process of cryptic exon inclusion during pre-mRNA to mRNA splicing. The variant frequency (VF) is defined by the formula: VF = reads variant / total reads event. (B) Volcano plot displaying significant splicing variants in iMN cells treated with TDP-43-ASO compared to the NT-ASO control. A total of 93 significant splicing variants were identified. (C) Volcano plot for iGN cells treated with TDP-43-ASO, revealing 192 significant splicing variants. (D) Scatter plot comparing the variant frequency (VF) of splicing changes in iGN and iMN cells after TDP-43-ASO treatment, highlighting variants that are significant in only iGN, only iMN, both, or neither. (E) UpSet plot showing the intersection of significant splicing variants (iGN spliced) and differentially expressed genes (iGN up, iGN down) within the iGN cell line. (F) UpSet plot showing the intersection of significant splicing variants (iMN spliced) and differentially expressed genes (iMN up, iMN down) within the iMN cell line. (G) Scatter plot comparing the variant frequency (VF) of splicing changes in iGN cells cultured with astrocytes or without after TDP-43-ASO treatment, highlighting splicing variants that are detected in co-culture (iGN+iAs) or monoculture (iGN) only, or in both/neither. (H) Bar graphs showing the VF for splice variants in STMN2, UNC13A, and HDGFL2 (n=2 for PBS, n=3 for NT-ASO and TDP-43 ASO).

To evaluate the potential influence of glial cells, which are also implicated in ALS pathology, we analyzed splicing changes in iGN co-cultured with iPSC-derived astrocytes (iAs). Most splicing alterations observed in iGN monocultures were also detected in the iGN-iAs co-cultures upon TDP-43 knockdown (Figure 3G, Table S5). Interestingly, the co-culture environment specifically facilitated the detection of a handful of additional cryptic events. Among these, we observed a cryptic event in Hepatoma-derived growth factor-related protein 2 (HDGFL2) (Figures 3G, H), an event that has previously been identified in the cerebrospinal fluid of ALS patients ^6^. The critical cryptic events in UNC13A and STMN2 genes were consistently detected in both iGN monoculture and iGN-iAs co-cultures (Figure 3H).

To validate our in vitro models against human disease, we compared the alternative splicing events in TDP-43-ASO-treated iGNs and iMNs with those described in post-mortem ALS and FTD patient tissues ^10^. We found that most cryptic splicing events identified in human post-mortem tissues were successfully recapitulated in our hiPSC-derived models (Figures 4A, C). Conversely, a few events detected in our in vitro systems, while absent in patient tissues, were reproducible across other human neuron in vitro models (Figures 4A, B), possibly representing a general in vitro TDP-43 depletion signature.

**Figure 4:**
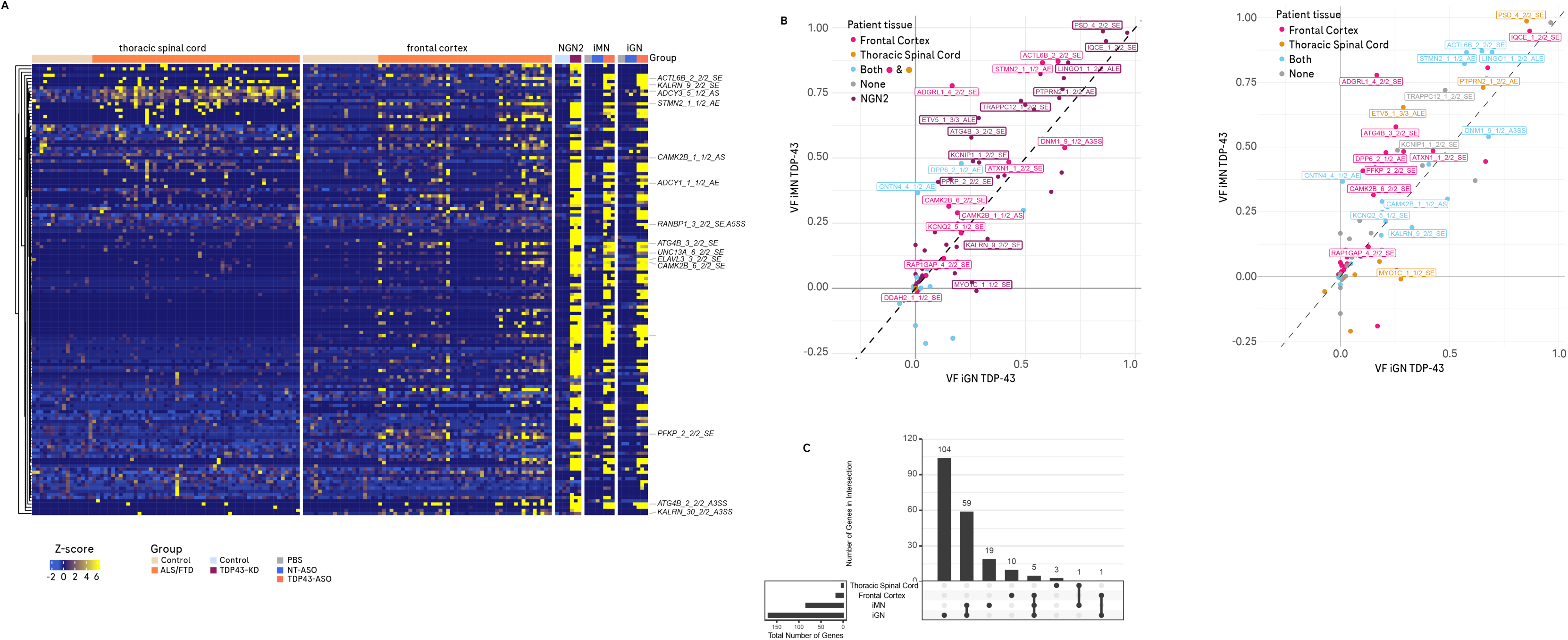
Splicing alterations in iGNs and iMNs relates to ALS/FTD patient data. (A) Heatmap showing the Z-Scores in splicing variants in human thoracic spinal cord and frontal cortex from ALS/FTD or control tissues compared to a public in vitro NGN2 dataset and our iMN and iGN cell lines. (B) Scatter plot comparing the variant frequency (VF) of splicing changes in iGN and iMN cells after TDP-43-ASO treatment, highlighting variants that are also significantly affected in patient frontal cortex tissue, thoracic spinal cord, both, neither or in vitro NGN2 neurons. (C) UpSet plot showing the intersection of affected genes in iGN or iMN in vitro cultures and patient frontal cortex and thoracic spinal cord tissue.

### TDP-43 knockdown induces a hyperexcitable phenotype

Neuronal hyperexcitability is a widely recognized functional hallmark of ALS neurons, both in patients and in in vitro models ^12–16^. To assess whether this phenotype is a functional consequence of TDP-43 knockdown, we performed Microelectrode Array (MEA) electrophysiology recordings in iGN/iAs or iMN/iAs co-cultures. At baseline, iGN and iMN cultures displayed fundamental differences in their neuronal network activity (Figures 5A, B and S2, S3), consistent with the core function of the main neurotransmitters in those cultures. After four or five weeks in culture, iGNs developed a highly organized and synchronous firing pattern with clear bursts (Figures 5A and S2), reflected by a high cross-correlation index (Figure S3). In contrast, iMNs exhibited only an asynchronous firing pattern, which persisted even after five weeks of culture, and showed a very low cross-correlation index (Figures 5B, S2 and S3).

**Figure 5:**
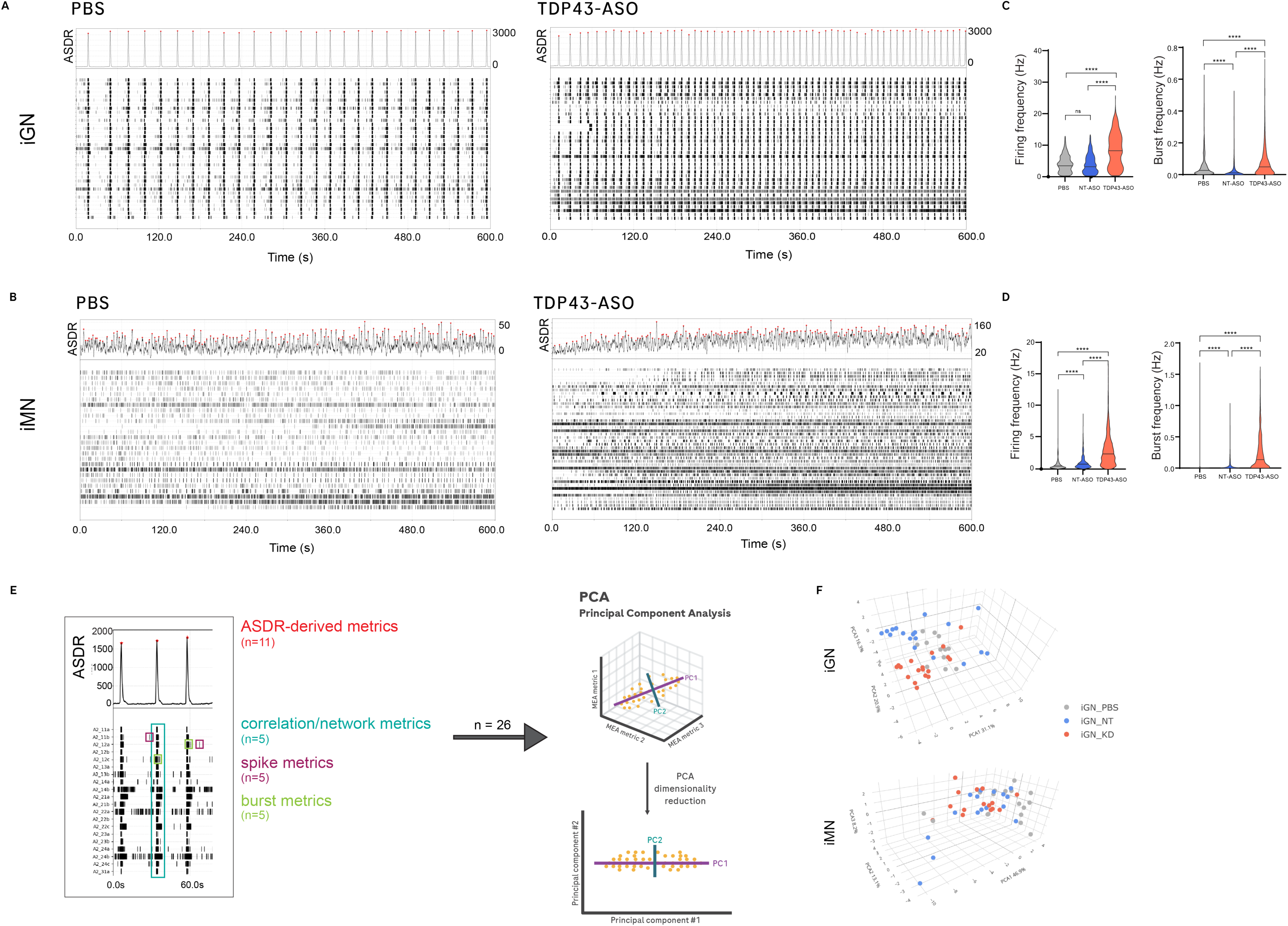
TDP-43 knockdown elicits a hyperexcitable phenotype in iNs. (A) Raster plots showing the spontaneous electrical activity of iGN cells treated with PBS (control) and TDP-43-ASO. The raster shows one neuron per line over a period of 600 seconds. The array-wide spike detection rate (ASDR) summarizes activity over all neurons and time. (B) Raster plots showing the spontaneous electrical activity of iMN cells treated with PBS (control) and TDP-43-ASO. The raster shows one neuron per line over a period of 600 seconds. The ASDR summarizes activity over all neurons and time. (C) Violin plots quantifying the firing frequency and burst frequency for iGN cells, comparing PBS, non-targeting (NT), and knockdown (KD) groups. Kruskal-Wallis with Dunn’s post-hoc (p**** < 0.0001). (D) Violin plots quantifying the firing frequency and burst frequency for iMN cells, comparing PBS, non-targeting (NT), and knockdown (KD) groups. Kruskal-Wallis with Dunn’s post-hoc (p**** < 0.0001). (E) Summary of the ASDR plot and its derived features, including the PCA analysis comprising 26 features. (F) 3D PCA plots visualizing the electrophysiological profiles of iGN and iMN cells, showing how the different treatments (PBS, NT, and KD) affect their overall electrical activity across 26 features.

Despite these differences in baseline network activity, TDP-43 knockdown resulted in a significant increase in the average firing rates (aFR) in both iGN and iMN cells (Figures 5C, D). We also observed increases in bursting frequency in both neuronal subtypes (Figures 5C, D). However, the nature of this increase differed: in iGNs, the increased burst frequency was accompanied by a higher number of spikes within bursts (Figure S3), whereas in iMNs, the average duration of bursts was higher (Figure S3). The lack of synchronized activity in iMNs persisted after TDP-43 loss-of-function, with the low cross-correlation index remaining unaffected (Figure S3). In iGNs, where synchronicity was high, TDP-43 knockdown also did not affect the cross-correlation index (Figure S3).

To further characterize the network activity changes, we analyzed additional features derived from the Array-wide Spike Detection Rate (ASDR) distribution ^19,20^. In iGNs treated with TDP-43-ASO, we observed an increase in the frequency of ASDR peaks and ASDR skew compared to both NT ASO and PBS controls (Figures 5, S3). This suggests that TDP-43 knockdown specifically enhances the population-level firing dynamics in neurons communicating via glutamatergic synapses. A Principal Component Analysis (PCA) combining 26 features extracted from the MEA recordings revealed a clear separation between iGNs treated with TDP-43-ASO and the control groups (PBS or NT-ASO), indicating a significant shift in the overall electrophysiological profile (Figure 5D). This separation was primarily driven by the ASDR-derived metrics. Interestingly, this divergence in the TDP-43-ASO iGN group could be observed as early as two weeks, before the networks achieved full synchronization (Figure S2). In contrast, the PCA performed on the iMN culture features did not show a clear separation of the TDP-43-ASO group from the controls (Figures 5F, S2), suggesting a less pronounced or qualitatively different effect on the network properties of lower motor neuron-like cells.

### TDP-43 knockdown does not affect neuronal susceptibility to glutamate excitotoxicity

The altered expression of key neuronal genes, such as CAMK2B, STMN2 and UNC13A, along with the functional impact of TDP-43 knockdown could negatively affect cell health and resilience to stress. To test this hypothesis, we first measured cell viability using a cell viability assay, neuronal morphology and the levels of Neurofilament light (NfL) in the supernatant, a marker of neuronal damage, after three or five weeks of culture. TDP-43-ASO treatment did not significantly impact cell viability after three or five weeks of culture (Figures 6 and S4). Similarly, neurite length and NfL levels were similar across all conditions (Figures 6 and S4), suggesting that TDP-43 knockdown had no significant effect on axonal degeneration and neuronal survival, despite the significant changes in gene expression and electrophysiological properties.

**Figure 6:**
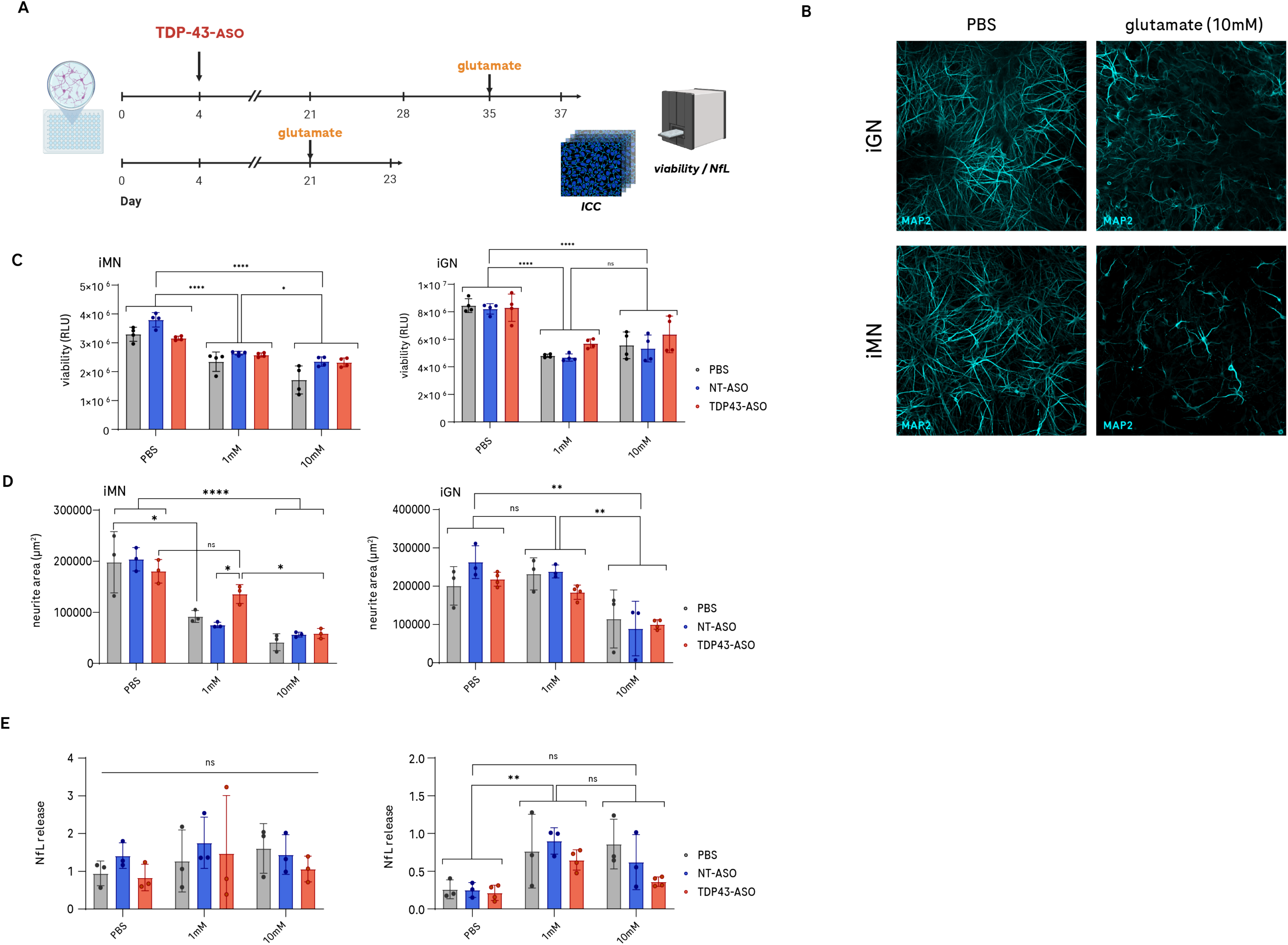
iN morphology and viability is affected by glutamate exposure, but not TDP-43 knockdown. (A) Experimental design for glutamate excitotoxicity experiments. (B) Representative images showing the morphology of neurites stained by anti-MAP2 in iGN and iMN cells under control (PBS) conditions and after treatment with 10mM glutamate for 48h. (C) Bar graphs demonstrating the effect of TDP-43-ASO knockdown on cell viability (relative luminescence units) for iGN and iMN cells with two concentrations (1mM, 10mM) of glutamate or PBS control. Cells were cultured in monoculture for 3 weeks (n=4), two-way ANOVA with Sídák post-hoc (p**** < 0.0001, p* < 0.05). (D) Bar graphs quantifying the dose-dependent effect of a 48h glutamate treatment (1mM, 10mM) on neurite length in both iGN and iMN cells, quantified from ICC MAP2 staining. Data points represent 3 independent experiments containing (n=3) samples. Two-way ANOVA with Sídák post-hoc (p**** < 0.0001, p** < 0.01, p* < 0.05). (E) Neurofilament light (NfL) release measured in both iGN and iMN culture supernatant after 5 weeks. Data points represent 3 independent experiments containing (n=3) samples. Two-way ANOVA with Sídák post-hoc (p** < 0.01).

We next investigated whether TDP-43 knockdown could alter the neuronal response to glutamate-induced excitotoxicity. We found that treatment with 5 or 10 mM glutamate for 48 hours caused morphological changes and reduced neurite length in both iGN and iMN cells and this reduction in neurite length was dose dependent (Figures 6B and S4). Glutamate significantly reduced iMN neurite length at concentrations as low as 1mM. In contrast, the effect of glutamate on iGN was only observed at doses of 5mM or higher (Figure S4). After treatment with 1 mM and 10 mM of glutamate for 48h in 21 days iMNs and iGNs cultures, we observed a significant reduction in both cell viability and neurite area (Figures 6C, D). However, the TDP-43-ASO knockdown cells showed no increased susceptibility to glutamate treatment, even at elevated doses, when compared to control groups (Figures 6C, D, E).

## Discussion

This study presents a comprehensive analysis of the molecular and functional consequences of TDP-43 loss-of-function in two distinct hiPSC-derived neuronal cell types: iGNs (upper motor neuron-like) and iMNs (lower motor neuron-like). Our findings validate the use of these models to study TDP-43 pathology and critically reveal cell-type-specific changes in gene expression, alternative splicing, and network excitability.

The number of differentially expressed genes and splicing variants identified in both iGNs and iMNs confirms the broad impact of TDP-43 dysfunction on RNA metabolism. However, the limited overlap between the DEGs of iGNs and iMNs underscores the importance of using relevant cellular models to uncover disease mechanisms specific to different neuronal populations. Whether these differences are related to specific roles of TDP-43 in gene expression regulation in glutamatergic and cholinergic neurons or could reflect some indirect effects of TDP-43 knockdown in those neuronal subtypes remains to be investigated. Nevertheless, our findings support previous studies highlighting cell-type-specific cryptic splicing signatures ^10,11^.

On the other hand, our results reveal a consistent downregulation and mis-splicing of established disease-relevant targets, notably STMN2 and UNC13A, in both glutamatergic iGNs and cholinergic iMNs. This strong, conserved molecular signature across two distinct neuronal subtypes is highly consistent with scientific literature. Loss of nuclear TDP-43 is known to cause the aberrant inclusion of cryptic exons, leading to the premature polyadenylation and subsequent decay of functional STMN2 mRNA ^7,8^. The resulting deficiency in functional STMN2 protein impairs microtubule dynamics, directly compromising motor neuron growth and repair—a defect strongly linked to ALS pathogenesis ^7^. Similarly, the mis-splicing of UNC13A leads to the loss of the functional, full-length protein ^18^. Given that UNC13A is a critical pre-synaptic protein mediating vesicle priming, its reduction impairs synaptic transmission and is suggested to contribute significantly to early synaptic dysfunction, a hallmark of ALS ^9^. The strong correlation in splicing changes for these core targets between iGNs and iMNs confirms that a fundamental, conserved TDP-43-mediated mechanism of RNA processing failure is at play, regardless of the neuronal subtype’s identity, thereby validating the relevance of our hiPSC models for studying core ALS pathology.

Functionally, our MEA electrophysiology analysis revealed that TDP-43 knockdown induces a hyperexcitable phenotype in both glutamatergic (iGNs) and motor (iMNs) neurons, characterized by increased firing and bursting. It also shows that TDP-43 knockdown in iGNs does not affect neural network synchronization. This contrasts with the findings reported by Keuss and colleagues (2024), who observed that TDP-43 depletion in human iPSC-derived neurons resulted in a severe reduction in synaptic transmission and a disordered and asynchronous pattern of network activity, a profile often associated with hypoactivity or impaired connectivity. This discrepancy likely stems from differences in the specific neuronal subtypes used and the knockdown levels achieved in the different models. Nevertheless, our observation of normal network activity even in the presence of a significant reduction in the expression of UNC13A in iGNs is intriguing and could suggest that this is not the main driver of the electrophysiological phenotype observed. Indeed, neuronal hyperexcitability (increased firing rate) may be independently driven by the mis-splicing of other critical targets involved in intrinsic membrane excitability, such as the KCNQ2 ion channel, which has been linked to TDP-43-dependent hyperexcitability in related studies ^21^.

Despite the robust and conserved molecular and functional alterations elicited by TDP-43 knockdown in both iGNs and iMNs, a striking paradox emerges: we observed no significant impact on overall neuronal survival, neurite length, or NfL release in the knockdown cultures. STMN2 is widely recognized as a crucial mediator of motor neuron growth and regeneration, and its depletion is expected to cause dendritic retraction and impaired neurite outgrowth ^7^. The lack of a morphological deficit in our models, despite five weeks of profound STMN2 depletion, suggests that hiPSC-derived neurons possess compensatory mechanisms or a sufficient residual level of STMN2 to maintain basal morphology in vitro. Alternatively, the function of STMN2 may be more critical for repair following injury or long-term maintenance in the complex in vivo environment, rather than for maintaining structure in a simplified, nutrient-rich culture.

More strikingly, TDP-43-depleted neurons did not exhibit increased susceptibility to glutamate-induced excitotoxicity. This suggests that glutamate and TDP-43 loss-of-function may not act synergistically to drive immediate death at the ∼50% knockdown levels achieved here. However, this does not preclude vulnerability to other stressors. Emerging evidence indicates that TDP-43 knockdown significantly sensitizes neurons to alternative challenges such as DNA damage (TDP-43 is essential for the non-homologous end joining repair pathway), oxidative stress and proteotoxic stress ^22,23^. Interestingly, the downregulation of ATG4B observed in our data suggests impaired autophagy ^24^, which may only become fatal when the neuron is challenged with the accumulation of misfolded protein aggregates.

In conclusion, our findings support the use of hiPSC-derived neuronal models to study TDP-43-related pathology and stress the critical importance of using distinct neuronal subtypes to uncover cell-type-specific disease mechanisms. Future research should focus on the specific molecular pathways that link the observed splicing and expression changes of critical targets like STMN2 and UNC13A to the hyperexcitable phenotype, which will be essential for developing targeted, cell-type-specific therapeutic strategies.

## Acknowledgments

The authors would like to thank the members of the Neuroscience and Rare Diseases department at Roche Pharma Research and Early Development for their technical assistance and insightful discussions throughout the course of this study. This research was supported solely by internal funding from F. Hoffmann-La Roche Ltd.; no external financial support or grants were received for this work.

## Data availability

The bulk RNA sequencing data, including gene expression counts and alternative splicing variant frequencies for pure iNs (iGNs and iMNs) and iN+iAs co-cultures at day 35, have been deposited in the Mendeley Data repository. This data can be accessed via DOI: 10.17632/rd8kt4brt5.2. Raw electrophysiological recordings (.nex files) documenting the longitudinal development of neuronal network activity from week 3 to week 8 are available from the corresponding author upon reasonable request. All differentially expressed genes (Tables S1 and S2) and significant splicing variants (Tables S3 and S4) identified in this study are included in the supplementary files of the manuscript.

## Material and Methods

### 1. Cell Culture and Maintenance

Human induced pluripotent stem cells (iPSC)-derived ioGlutamatergic Neurons and ioMotor Neurons were obtained from Bitbio (Cambridge, UK) and cultured according to the manufacturer’s instructions at 37°C in a humidified incubator with 5% CO2. For experiments involving co-culture, ioGlutamatergic and ioMotor Neurons were plated together with iCell Astrocytes (Fuji Film Cellular Dynamics, Inc.) at a ratio of 2:1. Cultures were maintained for a total of three to five weeks and media was exchanged every 2-3 days. For experiments involving microelectrode array (MEA) recordings, cultures were maintained for extended periods up to eight weeks, with regular media changes.

### 2. TDP43 Knockdown

To achieve knockdown of TDP43 cells were treated with 5 µM of a TDP43-targeting antisense oligonucleotide (ASO) on day 4 post-plating, added directly to the culture medium. Control conditions included treatment with a non-targeting ASO at the same concentration and a vehicle control (PBS). Following ASO administration, cultures were maintained under standard conditions for the remainder of the experiment.

### 3. Immunocytochemistry

Cells were fixed with 4% paraformaldehyde in PBS for 15 minutes at room temperature, followed by three washes with PBS. Cells were permeabilized and blocked with 0.1% Triton X-100 and 5% Bovine Serum Albumin (BSA) in PBS for 30 minutes at room temperature. Cultures were incubated overnight at 4°C with primary antibodies targeting neuronal markers and TDP43: anti-TDP43 (mouse monoclonal, Abnova, Cat. No. H00023435-M01, 1:250 dilution), anti-MAP2 (chicken polyclonal, Neuromix, Cat. No. CH22103, 1:1000 dilution), and anti-HuC/D (rabbit polyclonal, Abcam, Cat. No. ab184267, 1:500 dilution). Following primary antibody incubation, cells were washed three times with PBS. Fluorescently conjugated secondary antibodies, diluted in blocking buffer (100 µL per well), were then added: goat anti-mouse IgG (Alexa Fluor 488, Thermo Fisher Scientific, Cat. No. A32723, 1:800), goat anti-chicken IgG (Alexa Fluor 647, Thermo Fisher Scientific, Cat. No. A32933, 1:1000), and donkey anti-rabbit IgG (Alexa Fluor 568, Thermo Fisher Scientific, Cat. No. A10042, 1:250). Secondary antibodies were incubated for 2 hours at room temperature. After secondary antibody incubation, cells were washed once with 100 µL of PBS before counterstaining. Nuclei were stained with DAPI (1:5000 dilution in 0.1% Triton X-100 in PBS) for 3 minutes at room temperature. Finally, wells were washed three times with PBS and left in PBS until imaging on a Opera Phenix Plus System (Revvity, Inc) 40x or 20x objectives and analyzed with Harmony High-Content Imaging and Analysis Software (Revvity, Inc).

### 4. Viability and Neurofilament Light (NFL) Assays

Cellular viability was assessed using either the Promega CellTiter-Glo Luminescent Cell Viability Assay or the Promega RealTime-Glo MT Cell Viability Assay, according to the manufacturer’s instructions. Luminescence readings were taken using a TECAN Plate Reader and normalized to the untreated control. For experiments using glutamate treatment, L-Glutamic acid (SIGMA-ALDRICH) in solution was diluted in cell culture medium and added to the supernatant at appropriate concentrations. Neurofilament light (NfL) protein levels were quantified from the culture supernatant. Supernatants were collected at the end of experiments and stored at −80°C until analysis. NfL concentrations were measured using a kit from Uman Diagnostics (Umeå, Sweden) following the manufacturer’s protocol with a dilution of 1:1200.

### 5. TDP-43 quantification

Protein TDP-43 levels were measured in the cell lysates (diluted 1:50) using Human TDP43 ELISA Kit (Abcam), according to the manufacturer’s instructions. Absorbance was recorded on a TECAN Plate Reader and normalized to the total protein concentration measured by BCA (Thermo Scientific, Pierce BCA Protein Assay).

### 6. RNA Extraction and Bulk RNA Sequencing

Total RNA was extracted from frozen cell pellets of the iGN and iMN cells. Total RNA was quantified with Qubit RNA HS Assay Kit (Thermo Fisher Scientific). RNA integrity was assessed using a Fragment Analyzer. Library preparation was performed using polyA selection for mRNA species to remove ribosomal RNA, generating Illumina-compatible libraries. The libraries were sequenced on an Illumina NovaSeq 6000 system (v2.0.0) to generate 2×150 bp paired end reads, with an average of 50 million reads per sample.

### 7. Analysis of RNA Sequencing Data

Analysis was performed according to methods previously described ^10^. Briefly, reads were aligned to the human reference genome GRCh38 using STAR aligner (v2.7.1) with default parameters. Gene counts were calculated from the sorted and indexed BAM files using featureCounts (v2.0.6).

For the alternative splicing analysis, the SGSeq package was used to discover and quantify splicing events ^25^. The relative usage of each splice variant (variant frequency, VF) was calculated as a ratio of the junction reads supporting the start or end of the variant versus the sum of reads for all variants in a splice event. SGSeq analysis was performed in one run for all samples.

DEXSeq was used to test for differential usage of splice variants within an event ^26,27^. Statistical significance for differential splicing was set at a False Discovery Rate (FDR) of 5% and a ΔVF (change in Variant Frequency) of 0.1. Cryptic splices were specifically identified as significantly differentially spliced variants where the control group mean VF was 0.05 or less. Testing was performed separately within each condition (iGlutamatergic Neurons; iMotor Neurons; iGlutamatergic Neurons + iAstrocytes; iMotor Neurons + iAstrocytes) to optimize cryptic splice detection.

Differential gene expression analysis was performed using the DESeq2 R package (v1.36.0) ^28^. Genes with an adjusted p-value (FDR) less than 0.05 and an absolute log2 fold change greater than 1 were considered differentially expressed.

### 8. Microelectrode Array (MEA) Recordings

ioGlutamatergic Neurons and ioMotor Neurons, co-cultured with iCell Astrocytes, were plated onto AXION MEA 48-well plates (AXION BioSystems, Atlanta, GA, USA) and treated with either TDP43-targeting ASO or control ASOs as described previously. Cells were seeded at a density of 30’000 neurons and 15’000 astrocytes per well. Cultures were maintained in a humidified incubator at 37°C with 5% CO2, with regular medium changes every 2-3 days. Electrical activity of the neuronal networks was recorded for 10 minutes weekly from week 3 to week 8 using an AXION Maestro Pro Multi-well MEA System (AXION BioSystems). Raw data was acquired using Axion Neural Recorder software (v3.12.6) and saved as .nex files.

### 9. MEA Data Analysis

Raw .nex files from the MEA recordings were spike sorted using Plexon Offline Sorter (v4.4.0) with kMeans (sortDim=2, scanStart=2, scanEnd=7, scanStep=1, scanStat=PsF) algorithm. Spike detection was performed using a 5.5-standard deviation threshold. Sorted .nex files were processed and analyzed using custom Python script (patent application P60705-EP-1 titled “Neuronal activity characterization in multi-channel systems”).

The average firing rate (aFR) was calculated for all electrodes above a minimum firing threshold of 0.2 Hz. Network burst detection was performed using the Array-wide Spike Detection Rate (ASDR) (Wagenaar et al., 2006), to identify population-level bursts. A burst was defined as having a maximum interspike interval (ISI) of 100 ms and a minimum of 3 spikes to be considered a valid burst. Peaks in the ASDR were identified with a minimum peak distance of 1000 ms and a prominence of at least 25% of the data range.

### 10. Analysis and statistics

#### ICC

For the ICC experiments, nine fields-of-view were acquired from the center of each well to calculate a mean per well. To measure neurite length, the total area of MAP2 labeled neurites was measured for each well. The sum of each well was then plotted by condition for which statistical tests (one-way ANOVA followed by Dunnett’s post-hoc correction) were performed to determine differences between conditions.

For quantification of TDP-43 knockdown, data from three independent experiments, each with three replicate wells, were analyzed. TDP-43 intensity was quantified in the nuclear region (DAPI labeled) and a mean intensity per well was calculated. For each independent experiment a global mean for each condition was then determined for each experiment. This global mean was normalized to the PBS control within each independent experiment. The final analysis to identify significant differences between the global means of conditions was a one-way ANOVA followed by Dunnett’s post-hoc correction.

#### Viability Assay

Viability was assessed using a luminescence readout. The relative luminescence unit (RLU) was measured for each well, with three replicate wells per condition. Each well’s measurement was normalized to the mean of all PBS control wells to account for inter-experiment variability. Depending on the specific experimental design, a one- or two-way ANOVA with an appropriate correction for multiple comparisons was employed to determine statistical significance.

#### Neurofilament Light Chain (NfL)

For NfL measurements, the absorbance of each well was read at 450 nm. A standard curve was generated, and the NfL concentration (pg/mL) for each well was extrapolated from this curve. To facilitate comparison across experiments, the NfL concentration was normalized to the mean of the PBS control condition for each experiment. Statistical significance was then determined using a one- or two-way ANOVA with an appropriate correction for multiple comparisons.

#### MEA

For the MEA data, both spiking and burst frequencies per neuron were analyzed. Given the non-normal distribution of this data, a Kruskal-Wallis test was used, with Dunn’s post-hoc correction applied to identify significant pairwise differences. Additionally, Principal Component Analysis (PCA) was conducted to reduce the dimensionality of the data from 26 extracted features. The results of the PCA were visualized in a 3D plot, where data points were colored by their respective experimental conditions to illustrate the separation of groups.

## Supplementary Figure captions

**Figure S1:**
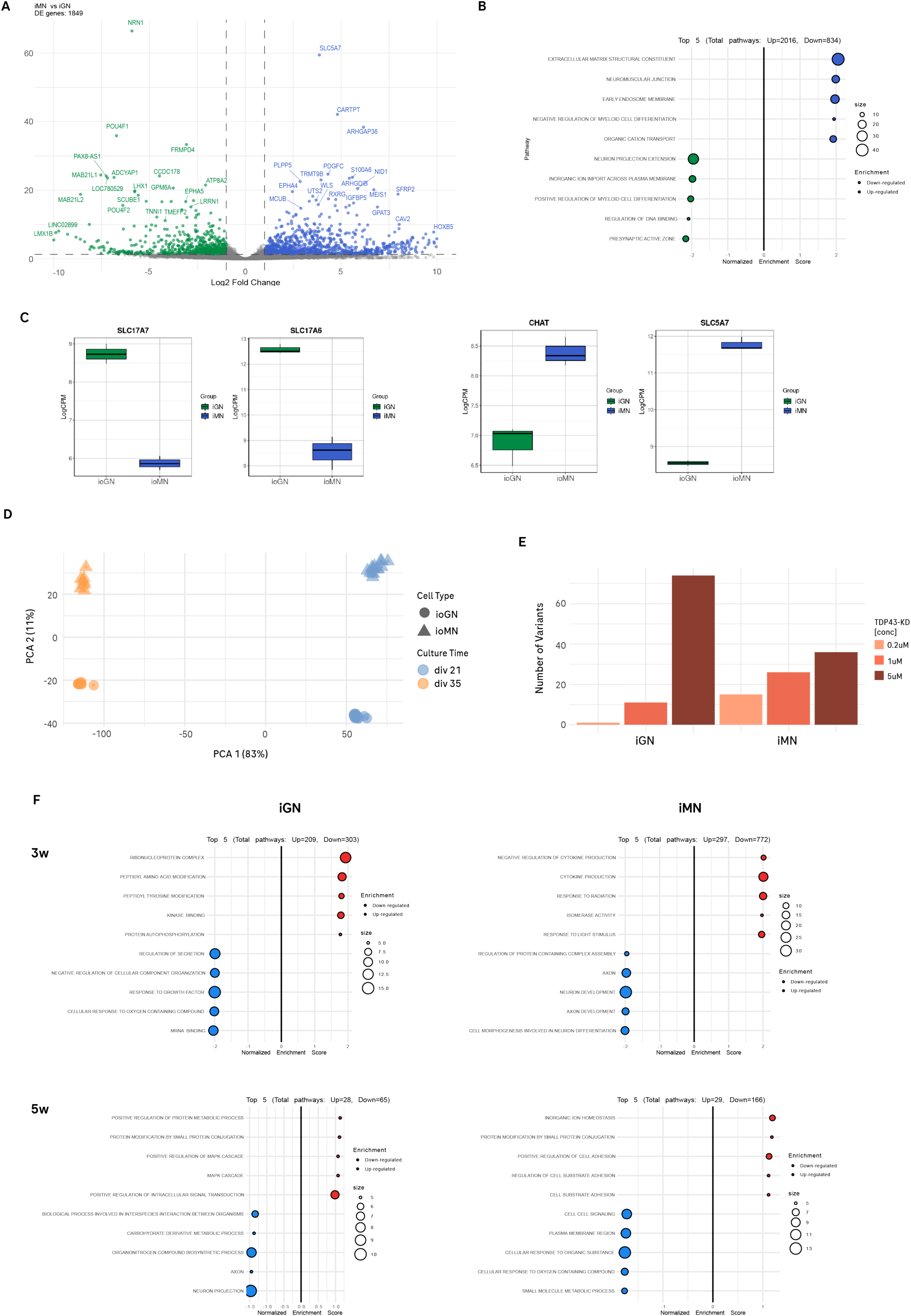
Molecular characterization og iGNs and iMNs used in the study. (A) A volcano plot showing 1,849 differentially expressed genes (DEGs) identified when comparing iMNs and iGNs. (B) Gene Set Enrichment Analysis (GSEA) of the top 5 upregulated and downregulated pathways in iMNs compared to iGNs. (C) Box plots of RNA-seq data confirm the identity of the models. iGNs show high expression of glutamatergic markers SLC17A7 (VGLUT1) and SLC17A6 (VGLUT2), whereas iMNs are characterized by the cholinergic markers CHAT and SLC5A7 (ChT). (D) Global transcriptomic profiles show distinct clustering of the two neuronal populations. PCA 1 (83% variance) primarily separates cells by cell type (iGN vs. iMN), while PCA 2 (11% variance) captures differences in culture time (day 21 vs. day 35). (E) Quantification of splicing variants following treatment with different doses of TDP-43 ASO. (F) Gene Set Enrichment Analysis (GSEA) of the top 5 upregulated and downregulated pathways in both cell types at day 21 (3 weeks) and day 35 (5 weeks) following TDP-43 depletion.

**Figure S2:**
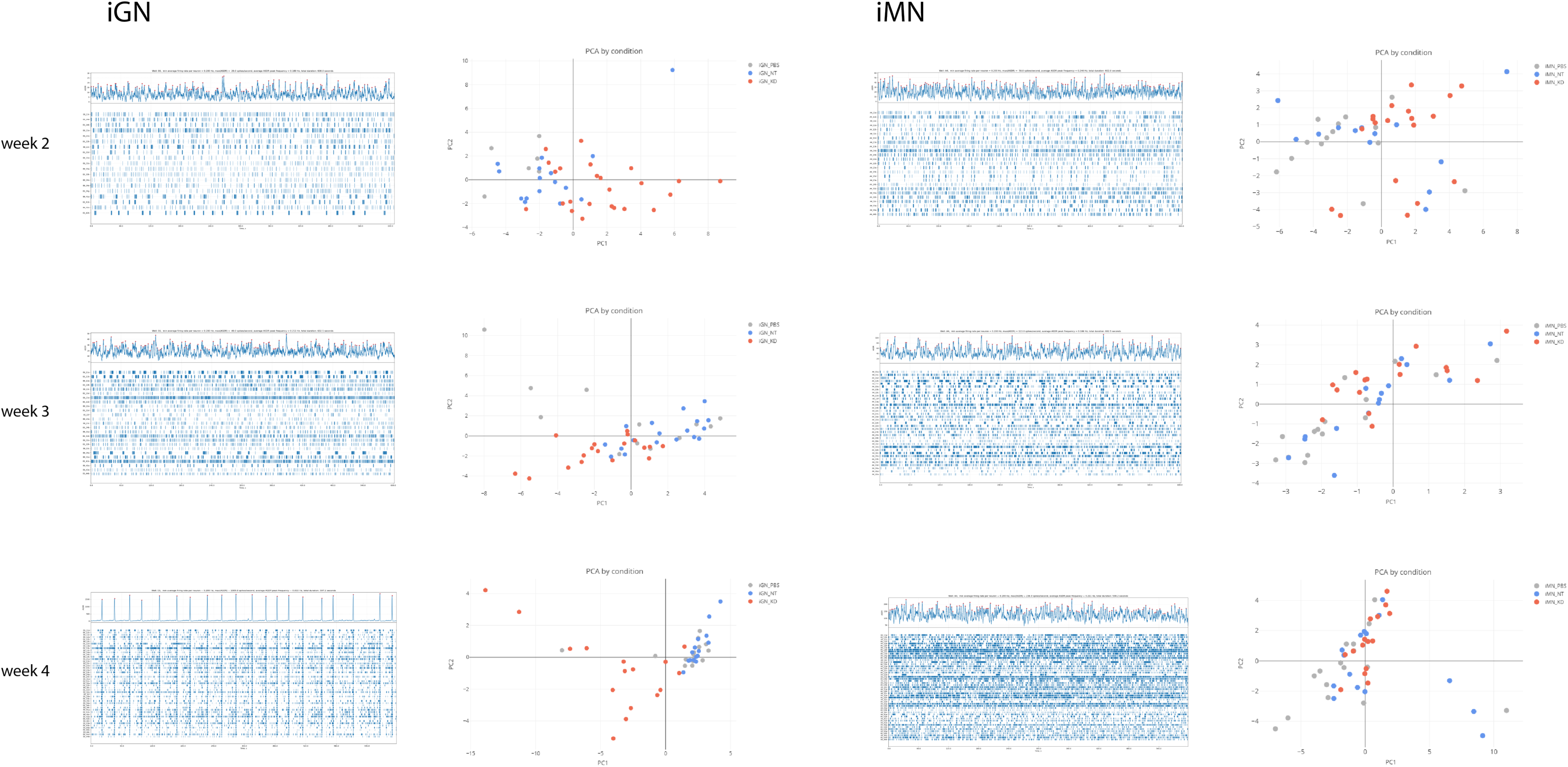
PCA based on eletrophysiological features distinguish iGNs trated with TDP-43 ASO from controls as early as after 2 weeks of culture. Raster plots (left panels) show representative 600-second recordings of spontaneous activity from week 2 to week 4 post-plating in iGN or iMN cultures. While, iGN networks progressively develop highly organized, synchronous firing patterns characterized by clear vertical alignments (network bursts) by week 4, iMN cultures maintain a predominantly asynchronous firing pattern throughout the developmental timeline. Principal Component Analysis (PCA) of using 26 extracted features (Right Columns) shows that a clear separation between the TDP-43-ASO group (red/pink) and control groups (PBS in blue, NT-ASO in grey) is detectable in iGNs as early as week 2, becoming more pronounced at week 4, when network synchronization is more prominent.

**Figure S3:**
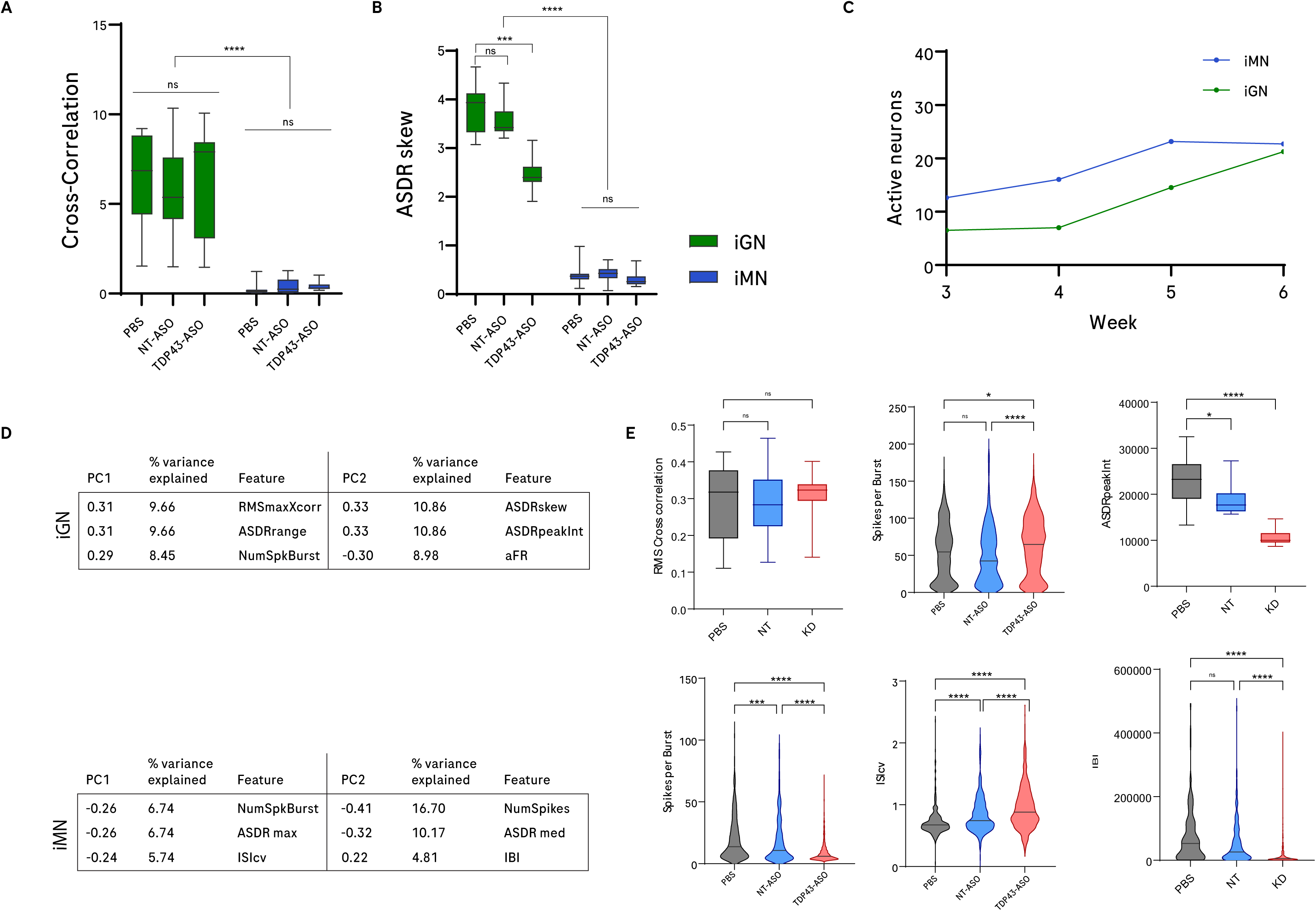
Detailed electrophysiological metrics and network synchronization analysis. (A) Box plot showing the cross-correlation (proxy of network synchrony) in iGNs and iMNs. Note that iGNs exhibit high baseline cross-correlation, which remains unaffected by TDP-43-ASO treatment, whereas iMNs display significantly lower baseline synchrony, also unchanged by knockdown. (B) Quantification of the Array-wide Spike Detection Rate (ASDR) distribution. iGNs treated with TDP-43-ASO show a significant decrease in ASDR skew compared to controls, reflecting enhanced population-level firing dynamics. (C) Longitudinal tracking showing the number of active electrodes from week 3 to week 6. (D) Tables listing the top features contributing to the first two principal components (PC1 and PC2) used in Figure 5. (E) Statistical comparison of selected features used in the PCA (Kruskal-Wallis with Dunn’s post-hoc; *p<0.05; ***p<0.001; ****p<0.0001).

**Figure S4:**
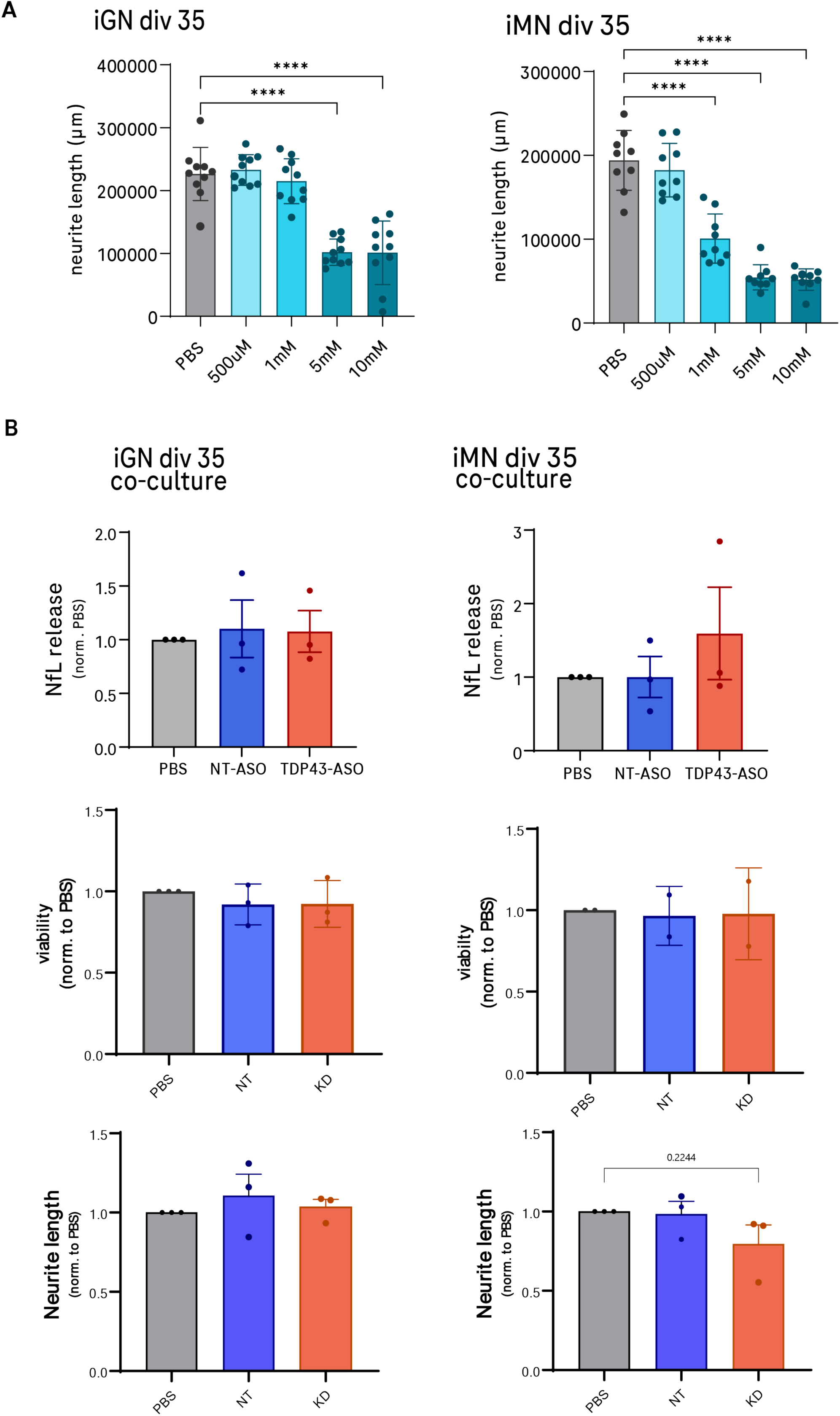
Dose-dependent response to glutamate-induced stress and evaluation of TDP-43 knockdown impact on neurodegeneration hallmarks. (A) Bar graphs quantify the effect of increasing glutamate concentrations (500 µM to 10 mM) on neurite length. iGNs show significant reduction in neurite length primarily at high doses of 5 mM or higher whereas iMNs exhibit higher sensitivity, with significant morphological deficits appearing at concentrations as low as 1 mM. (B) Quantification of NfL in the culture supernatant, cell viability and neurite length in 5 weeks cultures under basal conditions.

